# Adaptive surface sensing enables mammalian sperm navigation in complex environments

**DOI:** 10.1101/2025.11.16.688464

**Authors:** Ali Karimi, Xieergai Jiang, Alireza Abbaspourrad

## Abstract

During transit through the fallopian tube of the female reproductive tract, sperm encounter a structurally complex lattice-like environment characterized by confinement and curvature created by the folded epithelium of the fallopian tube. To replicate these conditions, we examined sperm scattering and migratory behavior in microfabricated obstacle lattices with varying spacing. We found a passive adaptive sensing mechanism that regulates sperm surface interactions in confined and curved environments. Specifically, scattering dynamics, including the collision angle, were impacted as sperm navigated the lattice, and surface interaction times were reduced under more confined conditions. A mathematical model was developed to describe sperm transport in these structured landscapes, predicting up to a nine-fold enhancement of diffusivity compared to free swimming sperm. Hyperactivated sperm displayed similar adaptive behavior within the lattice. We found that sperm dynamically adjust their navigation strategy in response to environmental geometry, allowing efficient migration despite confinement. More broadly, the study highlights how physical structure modulates microswimmer behavior and provides new insight into the physical processes that govern sperm navigation and the mechanisms underlying mammalian fertilization.

## Introduction

The environmental complexity of the features through which microswimmers move define their migratory behavior^1^. As an example, motile bacteria move faster in a colloidal suspension^2^ and their presence also causes the self-aggregation of passive particles^3^; however, the transport of the bacteria can be hindered ^4,5^ or enhanced^6^ by microporosity. Other biological microswimmers, including microalgae^7^, ciliates^8^, and nematodes^9^ exhibit similarly complex behaviors in structured environments. Despite its importance, there is a lack of understanding of how mammalian sperm interacts with a complex environment and how changing the features of the environment affects their migratory behavior.

Sperm motility is a principal determinant of mammalian fertilization^10^, whereas the oocyte is passively transported toward the fertilization site by the coordinated action of cilia and muscular contractions within the fallopian tube^11,12^. Sperm actively propel themselves through the tubal lumen using their flagella while continuously interacting with the epithelial surfaces^13–15^. These surfaces provide navigational cues that steer sperm toward the fertilization site^14,16–18^, impose selective pressures on the population^19,20^, and trigger essential biochemical signaling^21–23^. Their structural complexity, arising from the intricate folding of the epithelium, generates confined^24,20,25^ and curved^26^ surfaces whose degree varies along the length and radius of the fallopian tube^27,13^ (**Fig. 1a**). This interplay of curvature and confinement in such environments, which determines the dynamics of sperm interaction and migration, remains poorly understood.

**Figure 1.**
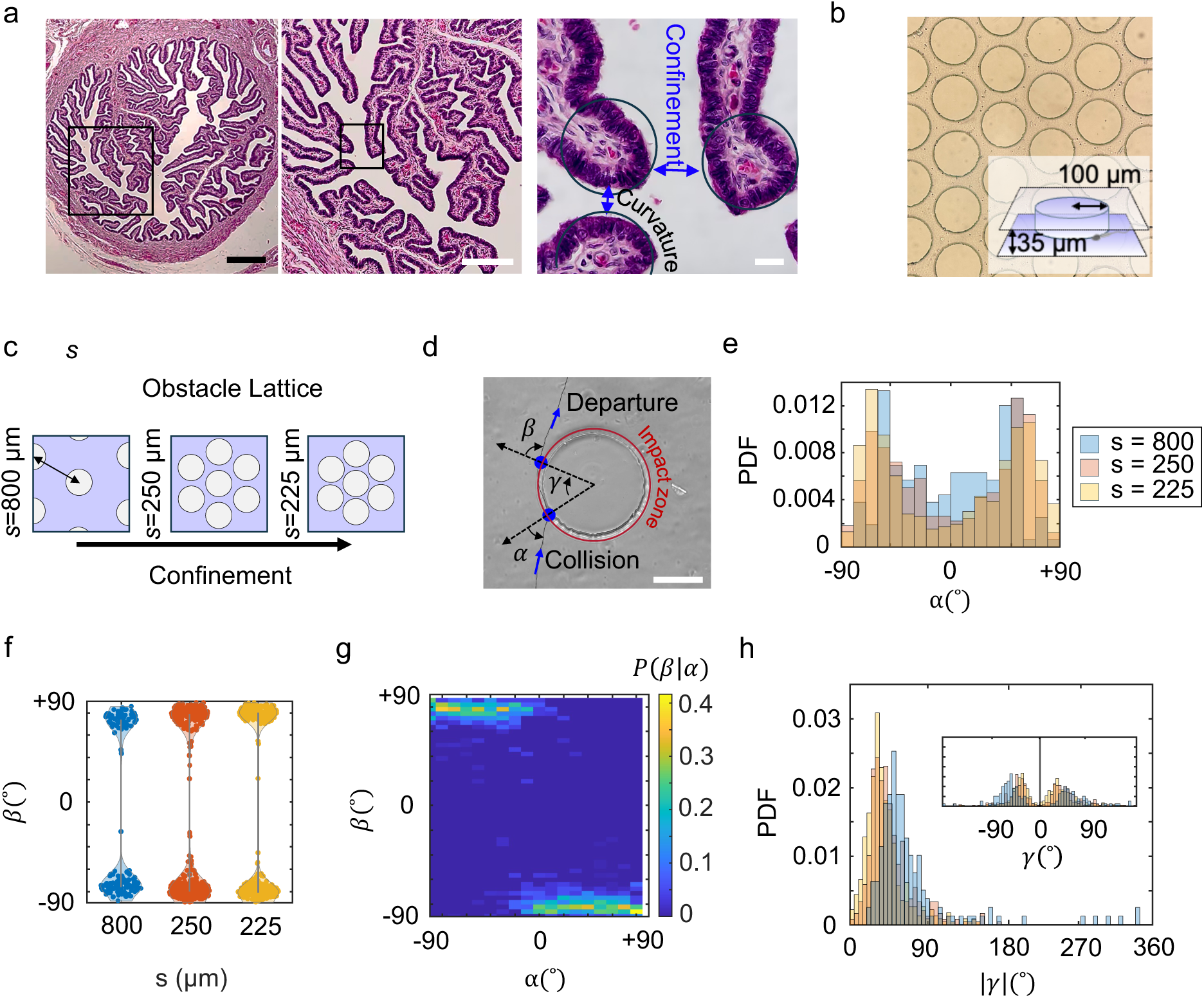
Sperm migration in the complex environment of the fallopian tube. (a) Cross-section of human fallopian tube showing curved and confined mucosal surfaces. Scale bar is 500 µm. Photos credit: Adolfo Sanchez-Blanco. (b) Bright-field image of the fabricated micro-obstacle array resembling the fallopian tube complexity. Inset shows the schematic of a single obstacle. (c) Variation of confinement by tuning the lattice spacing between obstacles. (d) Representative scattering event of a sperm with an isolated obstacle, illustrating definitions of scattering angles. (e) Probability distribution of collision angles (𝛼) within obstacle lattices. (f) Comparison of departure angle (𝛽) distributions for lattices with different spacings. (g) Conditional probability map P(β∣α), compiled from N=1117 interaction events across all lattice spacings. (h) Distributions of central angle |𝛾|. Inset: distributions of 𝛾.

Sperm are intermittently attracted to flat surfaces which results in sperm accumulation^28^ and two-dimensional beating in proximity to the surfaces^29^. When confined between two parallel surfaces, however, sperm exhibit a circular trajectory due to asymmetric beating^30,31^. The topology of the epithelial surfaces of the female reproductive tract (FRT) are more complex than simple flat surfaces (**Fig. 1a**) and exude significant influence over sperm migration. Specifically, the three-dimensional folding of epithelium creates highly convoluted geometry which alters the sperm-surface interactions and influences sperm guidance. For example, at junctions between two planes, sperm often follow the intersecting boundary until reaching a convex corner, where they scatter at sharp corners^32,16,33^ or follow the curvature if the corner radius exceeds ∼150 µm^16^. In contrast, at a concave corner, they tend to accumulate or become temporarily trapped^34–36^. Curvature is a defining feature of biological systems^37^, including the epithelial surfaces of the fallopian tube, where it governs how sperm hydrodynamically interact with their environment^26^.

In a curved concave boundary, the cells transition from a progressive, surface-aligned motility mode to an aggressive, surface-penetrating behavior as the local curvature increases^26^. If the epithelia of the fallopian tube is considered to be a continuous surface, then, by the Gauss–Bonnet theorem^38^, such concave (negative Gaussian curvature) must be balanced by convex (positive Gaussian curvature) regions to satisfy global geometric constraints. The interplay of convex and concave curvature, along with narrowing space (**Fig. 1a**) as sperm approach the fertilization site creates a complex topography of natural pathways navigated by sperm in the fallopian tube.

Here, we engineered a microfluidic device consisting of a microarray of circular obstacles (**Fig. 1b**). Similar designs were used previously to investigate other microswimmers such as bacteria^39,6^, microalgae^7^, ciliates^8^, and nematodes^9^, and now tailored to probe how bovine sperm scatter, interact with surfaces, and migrate. The circular shape and the closely packed spacing of the obstacle lattice are intended to reconstruct the curved and confined environment of the fallopian tube. We found that increasing inter-obstacle confinement alters sperm scattering and reduces the length of their interactions with surfaces when compared to less confined obstacles. Collision angle emerged as a key determinant of sperm exploration, providing abrupt directional changes upon obstacle encounter. Using a two-state Markov chain, we developed a computational model to predict the long-term effective diffusivity of sperm in the environment and observed enhanced transport in the presence of the obstacle array, potentially explaining the necessity of complex epithelial surfaces in FTR. Similar behavior was confirmed for hyperactivated sperm. These findings highlight the importance of environmental topology^40^ in shaping sperm interactions and migration, shedding light on the complex process of mammalian fertilization.

## Results and Discussion

We used circular obstacles with a radius of 100 µm located in the middle of a polydimethylsiloxane (PDMS) microfluidic chamber with a height of 35 µm fabricated by soft lithography^41^ (**Fig. 1b**). The chosen radius approximates the curvature of the folded structures of the FRT (20–150 µm)^26^. The obstacles were arranged in a hexagonal arrangement with lattice spacing *s* of 800, 250, and 225 µm, corresponding to gaps between obstacles of 600, 50, and 25 µm (**Fig. 1c**). Bovine (*Bos taurus*) sperm were used as a model for mammalian sperm in this study. We used standard Tyrode albumin lactate pyruvate solution (TALP)^42^ as the baseline medium; and sperm were maintained at 38.5°C (bovine body temperature) during the experiments. Video-microscopy was performed on a phase-contrast microscope with 10X objective lens; 30-s movies were acquired at 30 frames per second. We tracked the sperm while approaching and interacting with the obstacle in the standard TALP medium (**Fig. 1d**). Additional details on experimental setup and trajectory analysis are provided in **Methods**.

### Sperm scattering dynamics in structured environments

We measured collision angle (𝛼), departure angle (𝛽), and central angle (𝛾) for sperm scattering from obstacles. The collision angle distributions exhibit a bimodal pattern, with symmetric peaks at both positive and negative angles (**Fig. 1e**). The observed non-zero collision angle median, especially in the unconfined obstacle case (|𝛼$| ≈ 50°) is a unique feature not observed in progressively swimming active Brownian particles^43^. Sperm, however, swim in curved paths in proximity to flat surfaces due to hydrodynamic interactions^29^. We hypothesize that the chirality of this pre-collision path alters the collision angle distribution. To test this, we set up a coarse-grained model to predict the collision angle distribution for a timeless particle moving in a circular path of radius *r* diffusing toward and colliding with an obstacle of radius R (details of the collision angle model can be found in the **Supplementary Information Note 1)**. The model correctly predicts that the curvature of the path shifts the collision angle toward a non-zero median similar to experimental observations (**Fig. S4**). Note that the model neglected the variation in the intrinsic path curvature (1/*r* =10^−4^–10^−2^ µm^-1^) indicating the phenotypic variability in the population (**Fig. 4a**). Experimentally, the presence of neighboring obstacles (s = 250 and 225 µm) shifted the location of collision angle distribution modes to |𝛼| = 73° showing that sperm prefer gliding over head-on collisions. Sperm departed from the obstacles with the same mean angle〈|𝛽| 〉= 77° regardless of the confinement state of obstacles (**Fig. 1f**).

The conditional probability map P(𝛽|𝛼) shows a weak 𝛼–𝛽 correlation measured across N = 1117 scattering events indicating that sperm exit the surface close to tangent which is often regarded as slide-off boundary condition^43^ controlled mainly by steric interactions (**Fig. 1g**). Steric effects align the sperm with the solid surface upon collision and mediate their departure providing steric hindrance between the beating flagella and the surface^29,44^. The symmetry of the probability map indicates that short-range interactions and scattering are isotropic at the population level and within the geometrical setup. The majority of cells (> 89%) have a central angle less than 90° indicating that cell trappings are rare, and that sperm partially rotate on obstacles before departing (**Fig. 1h**). Confinement reduced the 𝛾 median from 59° in unconfined obstacles to 43° and 36° for lattice spacings of 225 and 250 µm, respectively (Mann–Whitney U test, *p* < 0.0001). The distribution of signed 𝛾 is symmetric around zero with slight dominance of counterclockwise rotations. Our observations of sperm scattering from a 100 µm radius obstacle are consistent with previous geometrical predictions, which suggest that boundary following (entrapment) in sperm occurs when the obstacle’s curvature radius exceeds 150 µm exclusively^6^.

The probability density map shows a dependence of the central angle on collision angle (**Fig. 2a**). The central angle has an inverse correlation with the collision angle; sperm grazing the obstacles stay a shorter angular distance on the surface compared to head-on collisions (**Fig. 2b**). Linear correlation with proportionality constant of unity was observed when 𝛼 > 45°. We also measured the equivalent interaction distance (*l)* as *l* = 2𝜋R |𝛾| /180 and found that sperm traversed a distance equal to their cell body on the obstacle surface before departing (**Fig. S5**).

**Figure 2.**
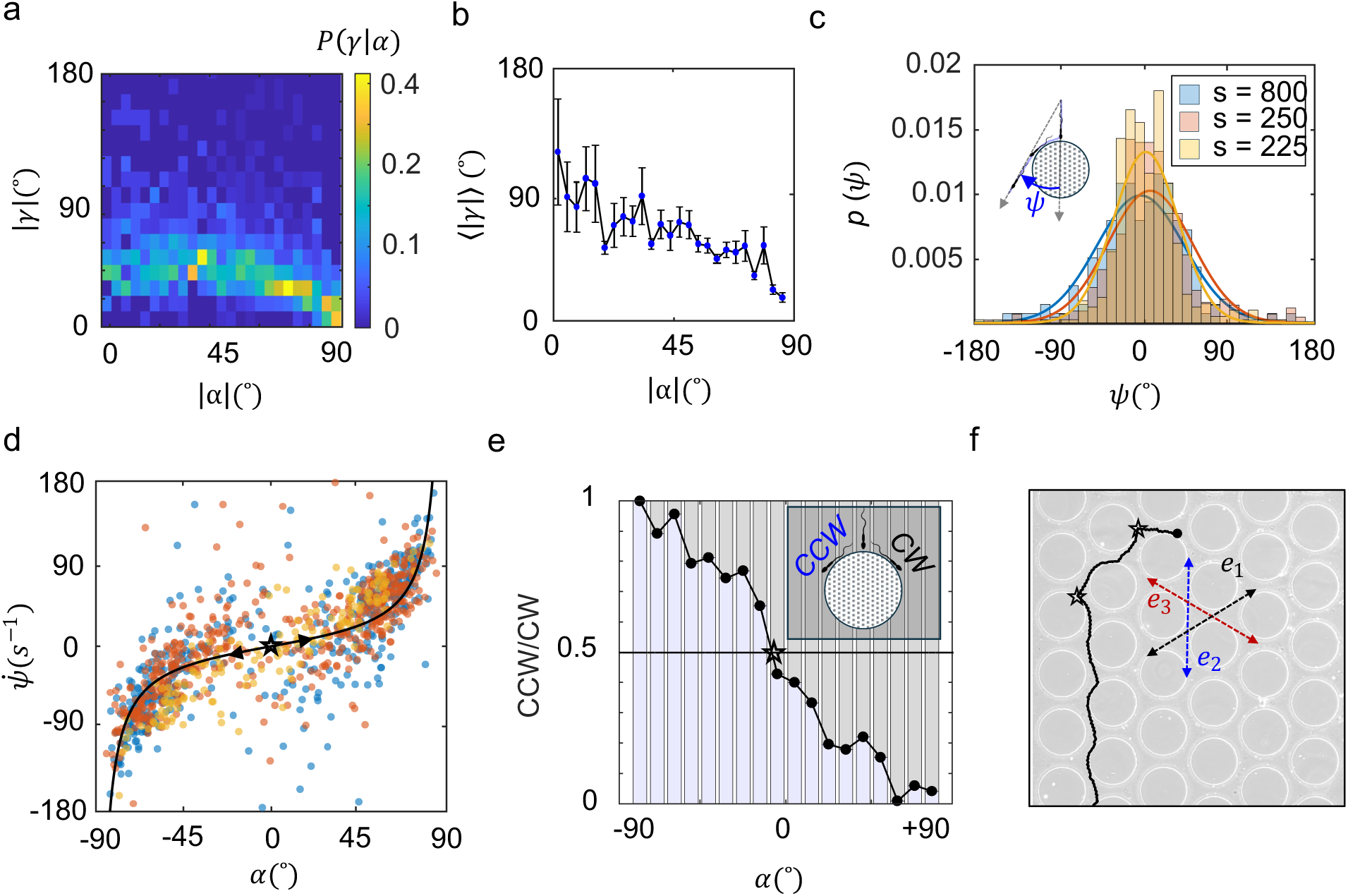
Sperm reorientation during interactions with obstacles. (a) Conditional probability map of |𝛾| and |𝛼|. The plot is compiled from N=1117 interaction events across all lattice spacings. (b) Mean central angle 〈|𝛾|〉 versus collision angle |𝛼|, where |𝛼| is taken as the center value of each bin, of width 5.5°. (c) Comparing the net reorientation 𝜓 distributions for different lattice spacings. Inset: sketch to define 𝜓. (d) Correlation between reorientation rate 𝜓̇ and 𝛼 color code with respect to lattice spacing. Solid line is the fit to the function 𝜓̇ = k tan 𝛼, with k being a constant. Star shows the metastable collision angle leading to a stochastic counterclockwise (CCW) or clockwise (CW) rotations. (e) proportion of CCW rotation in each collision angle. The dots show the average of each bin of width 10°. (f) Example trajectory of sperm exploring the 250 µm lattice with head-on collision indicated as stars and main lattice directions as e_1_–e_3_.

We define the net reorientation 𝜓 = 𝛼+𝛽-𝛾, which quantifies the change in sperm swimming direction resulting from an encounter with an obstacle within the lattice. The distribution of 𝜓 for the unconfined obstacle (*s* = 800 µm) is a Gaussian defined by p (𝜓) 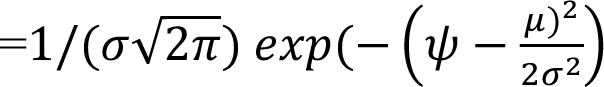 with a mean at µ∼0 and a standard deviation of 𝜎 = 46° (**Fig. 2c**).

Similar results were obtained for the confined obstacles with reduced standard deviations to 44° for *s* = 250 µm and 34° for *s* = 225 µm, indicating that sperm in the compact lattice undergoes fewer reorientations. We incorporated the reorientation time scale by defining the reorientation rate as 𝜓̇ = 𝜓/𝜏_i_ where 𝜏_i_ is the duration of interaction (collision to departure). The dependence of 𝜓̇ on the collision angle α is shown in **Fig. 2d**. Despite the scatter in the data across different lattice spacings, the results follow an overall trend characterized by vertical asymptotes at 𝛼 = ±90° and a plateau near zero. Analysis indicates that the data are well described by a simple tangent relationship 𝜓̇ (𝛼)= *k* tan(𝛼), where *k* = 23 s^−1^ is a constant reflecting the reorientation torque exerted by the obstacle together with other factors influencing scattering^44^.

A special case occurs when sperm hit the obstacle with 𝛼 ∼0° which results in a metastable condition where sperm remains still on the obstacle due to active normal propulsion against the wall until a small perturbation, either due to internal noise or external flagellum interactions with other obstacles, rotates the sperm head either clockwise (𝜓̇ <0) or counterclockwise (positive 𝜓̇ >0) (**Fig. 2d**). Measurement of the counterclockwise (CCW)/clockwise (CW) portion of the rotations of different collision angles showed that at 𝛼 ∼0°, there is an equal chance of rotating in either direction, CCW or CW (**Fig. 2e**), confirmed by high variation of data near this point (**Fig. 2d**). The resultant stochasticity of the rotation sign selection at 𝛼 ∼ 0° helps the sperm to explore the lattice between the three axes, e_1_, e_2_, and e_3_; for example, a sperm exploring a lattice with spacing of 250 µm might undergo two head-on collisions that shift its migration direction from e_3_ to e_1_ and then e_2_ (**Fig. 2f**).

### Confinement reduces interaction duration

We investigated the microscopic interactions of sperm with obstacles in confined lattices (s = 250 and 225 µm). While navigating the lattice, sperm transiently contact each obstacle before departing and moving toward the next (**Fig. 3a**). The distribution of interaction times is shown as survival curves S (𝜏_i_ > t), that is the probability of observing a contact time 𝜏_i_ greater than *t* (**Fig. 3b**). The distribution is composed of a deterministic time part, where S (𝜏_i_ > t) ∼1, and a stochastic part which the probability decays exponentially S (𝜏_i_ > t) ∝ exp(-k_D_ t), where k_D_ is the departure rate. The deterministic part of the contact time is required for all cells and is predominately the time required for sperm to locally align itself with the surface upon incidence.

**Figure 3.**
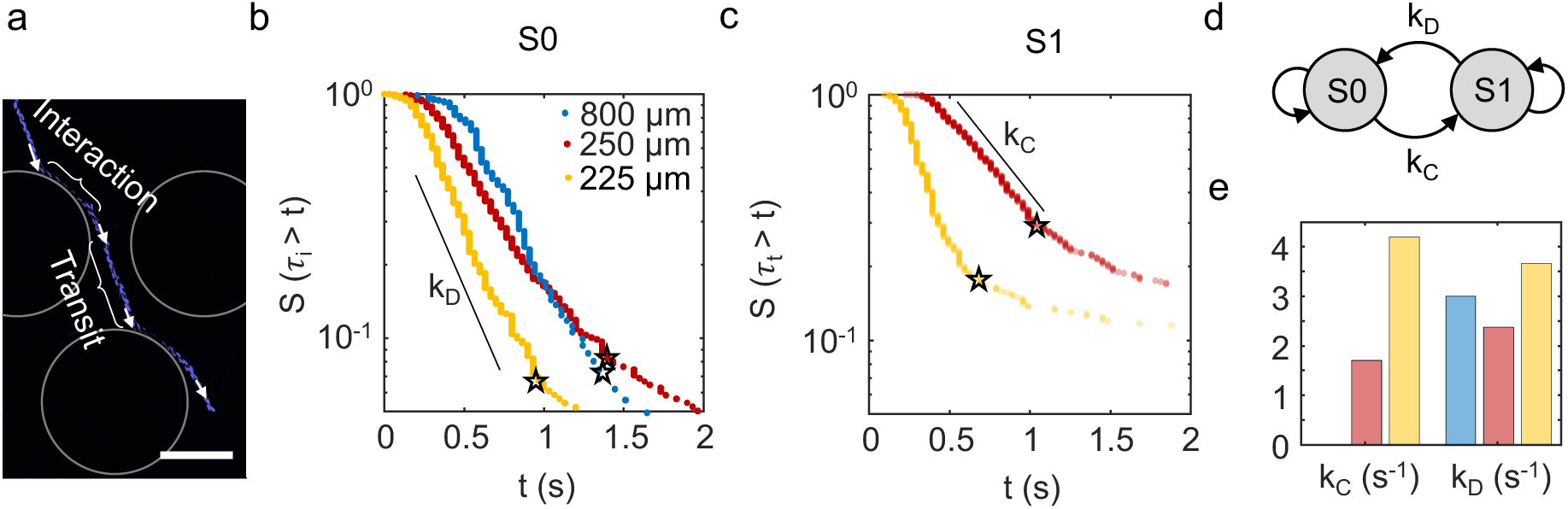
Duration of interaction. (a) An example of a spermatozoa interacting with obstacles within 250 µm spacing. The image is obtained from overlapping frames of sperm motion and subtracting the background from the image. Scale bar: 100 µm. Survival curves of (b) interaction and (c) transit durations. Stars indicate the transition to secondary exponential decays. (d) The resulting Markov chain model describing the sperm behavioral states in the obstacle lattice. (e) Collision (k_C_) and departure (k_D_) rates are estimated from the slopes in survival curves for lattices with spacings 800, 250, and 225 µm.

There is a secondary exponential segment where the decay rate is lower than the main exponential decay (**Fig. 3b** and **3c**, stars indicate the transition points). When compared between three spacings, we found similar deterministic and secondary exponential decay both in interaction (𝜏_i_) and transit (𝜏_t_) curves. We assign the secondary exponent to the phenotypic variation of cells which can have higher intrinsic chirality prolonging the interaction duration^45^. Therefore, we neglect the deterministic and secondary exponential parts in the distributions and investigate the motion in each state. Statistics of the stochastic part suggest that the sperm departure follows a Poisson process. This is qualitatively consistent with the random rotation of sperm flagellar waves relative to that of the head due to rotational Brownian motion which causes the sperm departure^29,34^. Next, we measured the duration of transit time while sperm jumped between obstacles and obtained a similar composition (a deterministic part and two exponential parts) (**Fig. 3c**). The deterministic part in 𝜏_t_ corresponds to the minimum distance between two jumps which we see extends for the lattice with smaller spacing. Using a similar approach as 𝜏_i_, we obtained the collision rate k_C_ which is the frequency of collisions within the lattice.

Since the interaction and transit phases are independent stochastic processes, the sperm behavior in a lattice of obstacles can be modeled by a simple two-state continuous time Markov chain (CTMC) (**Fig. 3d**) with master equations:

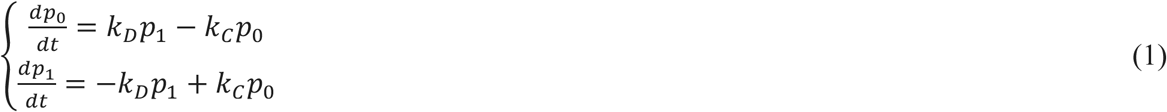

where p_0_ and p_1_ are the probabilities of finding a sperm in states 0 (transit, free space swimming) and 1 (interaction). Measurement showed the increase of k_D_ and k_C_ when the lattice spacing reduced from 250 µm to 225 µm (**Fig. 3e**). Although we do not have experimental measurements of k_C_ for the 800 µm spacing (unconfined obstacle), we can estimate it to be close to zero because of the large inter-obstacle distance. The steady-state probability of finding a sperm on the obstacle surface is given by 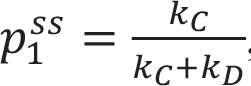, and we found that it decreases from 58% in the 250 µm lattice to 46% in the 225 µm lattice. We attribute this observation to near-field effects of adjacent obstacles on hydrodynamic forces^46,47^ (**Fig. S6**); the proximity of neighboring obstacle surfaces perturbs the flow field around a swimming sperm^48^ generating an imbalance in hydrodynamic forces that actively drives its detachment from the obstacle surface and attraction to the successive obstacle^49^. Another contributing factor is that the reduced mean free path in the confined lattice^50^ limits the time sperm spend in each state.

### Diffusive behavior

The microscale interactions with obstacles influence the macroscale transport of sperm within obstacle lattices and free space swimming (**Fig. 4a**). The short-range directed motion of sperm while interacting with obstacles was measured as the instantaneous orientation averaged over 1 s of each trajectory to obtain 〈𝜃〉 (**Fig. 4b**). In free space, only two broad peaks are observed which can be attributed to the bias caused by the cells diffusing from the inlet port.

**Figure 4.**
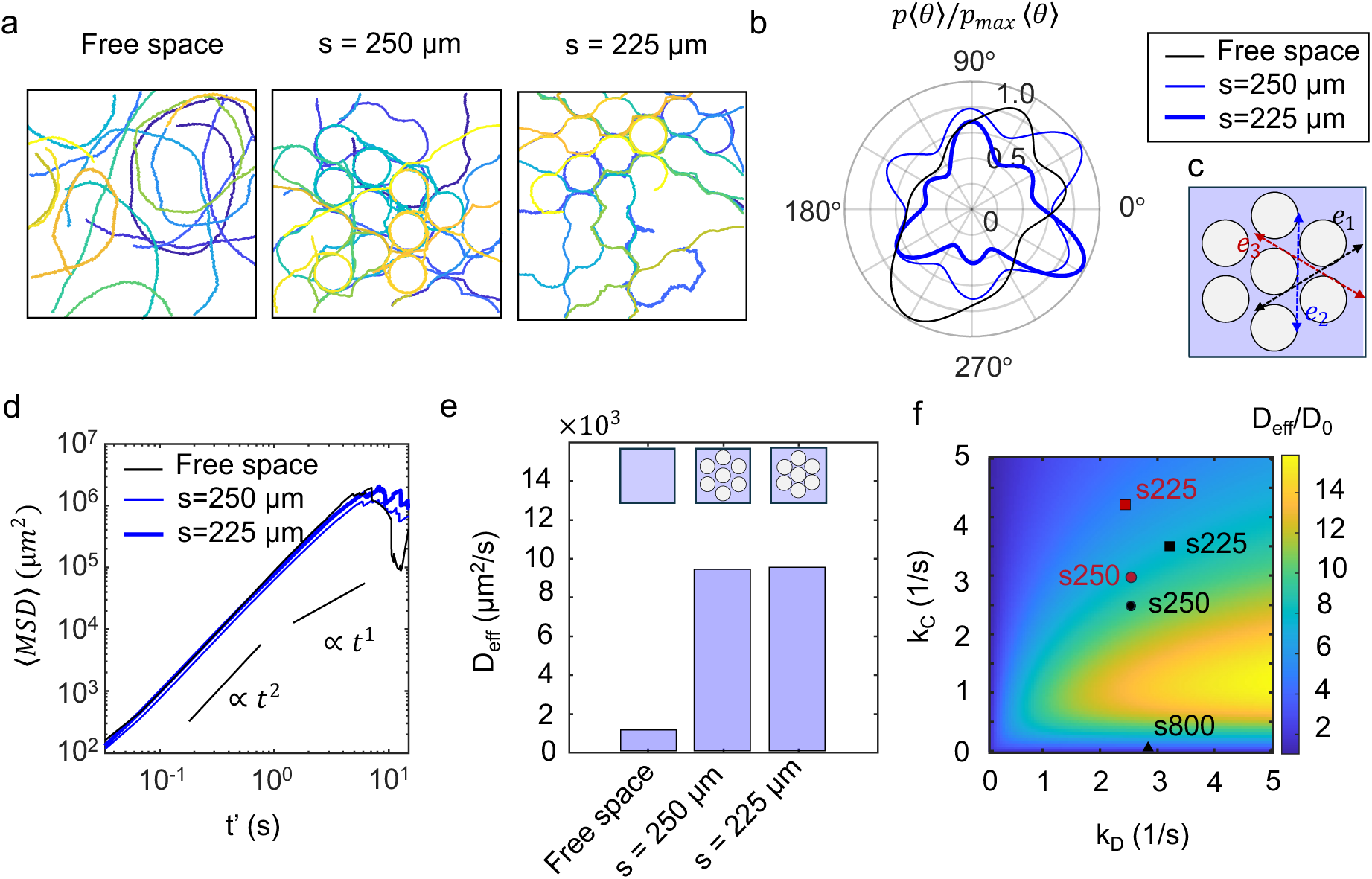
Diffusive behavior of sperm. (a) Example trajectories of sperm in free space and confined lattices selected randomly from the trajectories. (b) Normalized polar probability density of sperm swimming directions in lattices with spacings 250 and 225 µm and in free space. Instantaneous orientation vectors are averaged over 1 s to get〈𝜃〉. (c) Schematic showing the principal lattice directions. (d) Experimentally determined ensemble averaged mean squared displacement 〈MSD〉 over lag time *t’*. (e) Effective diffusivity of sperm transporting in free space and lattices obtained from Markov chain modeling. (f) The value of D_eff_ /D_0_ color-coded as a function of collision and departure rates, k_C_ and k_D_. Black and red markers correspond to experiments in TALP and TALP + 5mM caffeine, respectively.

Periodic interactions, observed in confined lattices, introduced additional peaks in the orientation distribution indicating the short-range directed motion in the lattices. Both obstacle lattices rectified the sperm motion by reducing the degree of freedom to 6 main directions (**Fig. 4c**). The peaks are exactly 60° apart from each other which is mapped to the main six lattice directions showing free pathways within the lattice. The magnitude of the three peaks is less pronounced when the confinement increases to 225 µm and we see three sharper equally spaced peaks.

Consistent with the increased probability of near-zero net reorientations, sperm in more compact lattices undergo less reorientations due to obstacle scattering which results in pronounced peaks (**Fig. 4**). The increased packing of obstacles causes directional locking^43,51^, which helps the sperm maintain their swimming direction while favoring gliding (|𝛼| closer to 90°) over obstacles.

The ensemble-averaged mean squared displacement 〈𝑀𝑆𝐷〉 = 〈𝑟(𝑡 + 𝑡′) − 𝑟(𝑡)〉#, where *r* is the location vector, showed ballistic movement for t’< 5 s (〈*MSD*〉∝ t’^2^) and then randomize their swimming direction showing diffusive behavior (〈*MSD*〉∝ t’); and then the curve starts to oscillate which makes analysis of it impossible (**Fig. 4d**). The transition point from ballistic to diffusive region is characterized by a crossover time which is the time the cells swim progressively before randomizing their direction. Cells in 250 µm lattice have a smaller crossover time (∼2 s) compared to the free space swimming cells (∼4 s) highlighting the role of stochastic scattering from obstacles. Similar behavior is reported for bacteria swimming in porous environments^39,52^. Increasing the lattice confinement extended this crossover time to ∼5 s showing a more persistent motion in the compact lattice. This is consistent with the higher directionality observed in instantaneous directions distribution (see **Fig. 4b**).

To investigate the long-term behavior of cells within the lattice requires sophisticated experimental setups and advanced computational tools^39^;therefore, we adopted an approach previously developed to characterize bacterial transport near surfaces^53^ and formulated a mathematical model grounded in the two-state CTMC framework. During the transition between obstacles (state 0), due to hydrodynamic interaction with the surface^29,54^, sperm follow a curved pathway with an angular velocity *𝛺_0_* and linear velocity *V_0_* along a time-dependent orientation angle *θ*(*t*) with respect to the *x*-axis. Sperm transitions between free-swimming and interaction states involve a net change in the moving direction *θ-θ’* (*θ’* is the orientation after interaction). The probability density of finding a sperm in transit or interaction states is given by:

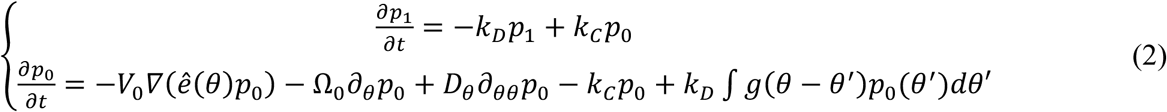

where *D_θ_* is the rotational diffusion coefficient and g(*θ*-*θ’*) is the reorientation kernel. This kernel is a function of net reorientation 𝜓 which is provided for the different spacings of the lattice in **Fig. 2c**. With the above model setting, we obtain an analytical expression for the effective diffusivity of chiral sperm in an obstacle lattice

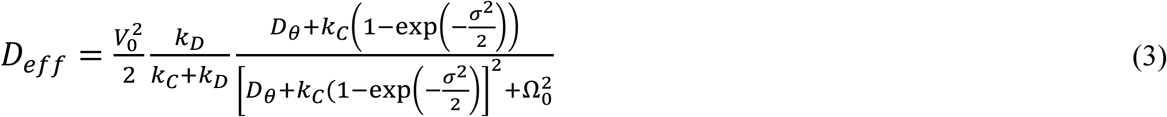

where *D_θ_* is the rotational diffusion of sperm head oscillation (details regarding the *D_eff_* derivation are provided in **Supplementary Information Note 3)**. 𝜎 was obtained from the standard deviation of the net angular reorientation distributions (**Fig. 2c**). Note that 𝜎 must be converted to radian to be able to use it in equation (3) Other parameters are obtained from the trajectories (N = 20) of free-swimming sperm: *D_θ_* = 0.01 rad^2^/s, *V_0_* = 186 µm/s, and *𝛺_0_* = 0.39 rad/s (see **Supplementary Information** and **Fig. S7**). For free-space unobstructed swimming of sperm (k_C_ = 0), equation (3) simplifies to

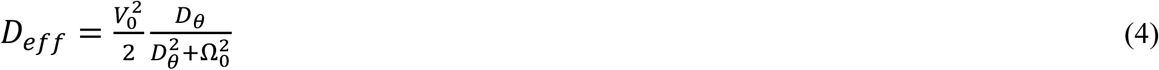

which is similar to the expression previously obtained for a chiral active particle^51^. Using the mean values, we obtain an effective diffusivity of 1.1྾10^3^ µm^2^/s for free swimming sperm, then for obstructed swimming (in obstacle lattice), 9.7྾10^3^ µm^2^/s for lattice with spacing *s* = 250 µm, and 9.9྾10^3^ µm^2^/s for *s* = 225 µm. The obstacle lattice increased the effective diffusion by 8.6-fold compared to free space swimming; further confinement (*s* = 225 µm) slightly enhanced it to 8.9-fold increase (**Fig. 4e**). Effective diffusion is influenced by k_D_ and k_C_; tuning the confinement modulates the two transition rates and results in a super-diffusion region where sperm are up to 7-fold more diffusive than free space (**Fig. 4f**). Increasing the collision rate, by decreasing obstacle distances, will reduce effective diffusivity due to the higher reorientation variation caused by collisions. We also can add the *s* = 800 µm to the points (k_C_∼0) where spacing is effectively large and collisions are rare. The values of the effective diffusivities increased in the complex environment, with negligible difference between the two confinements.

### Effect of hyperactivation

Hyperactivation is a specialized mode of motility characterized by high-amplitude, asymmetric flagellar beating that enables sperm to generate nonlinear motion^55,56^. In vitro, hyperactivation can be triggered by pharmacological agents or physiological cues^55^. We supplemented the TALP medium with caffeine at 5 mM to hyperactivate the cells^55^. Scattering behavior of sperm remained intact with hyperactivation (**Fig. S8**). A similar k-value was obtained from the 𝜓^.–𝛼 curve indicating similarity of overall behavior in lattice (**Fig. S9**).

Although not statistically significant (p > 0.05), the angular velocity and rotational diffusion coefficient increased in hyperactivated sperm swimming in free space, exhibiting higher intrinsic path curvature and lateral head oscillations (**Fig. S10**). Hyperactivated sperm had similar k_C_ in 225 µm lattice, however, this parameter increased in the 250 µm lattice (**Fig. S11**). The departure rate underwent a reduction from 3.2 for normal sperm to 2.4 for hyperactivated sperm in 225 µm lattice. In the more confined lattice, hyperactivated sperm spent more time on the obstacle compared to normal sperm. The ratio *D_eff_/D_0_* for hyperactivated sperm was 2.57 and 2.1 (for 250 and 225 µm spacings respectively), which is a quarter of the value for normal sperm. Therefore, hyperactivation magnified the effect of environmental complexity compared to free swimming (**Fig. 4f**). Overall, diffusivity increased with geometric complexity introduced by obstacles, whereas spacing, despite altering short-scale interaction and transit durations, had minimal effect on migratory behavior.

Since Antonie van Leewenhoek first observed sperm motility under the microscope in 1677^57^, numerous studies have sought to unravel sperm behavior, particularly within environments that mimic the female reproductive tract. However, it still remains unclear precisely how sperm migrate in a complex environment particularly with curved and confined surfaces. Using a hexagonal array of circular obstacles with different lattice spacings, we first quantified the scattering events using collision angle, departure angle, and central angles. We observed that the interaction of the sperm with the obstacles is controlled by steric effects^44^ and oblique angle collisions dominate due to chirality and the rectifying effect of the lattice regardless of confinement status. Despite being rare, head-on collisions are critical, as they enable sperm to explore the lattice through abrupt rotation change upon collision.

The duration of sperm–obstacle interaction is controlled by the stochastic Poisson process. Higher confinement, corresponding to a gap size of 25 µm, reduced the mean residence time on obstacles by 25% compared to less confined cases (gap size of 50 µm and 600 µm). Regarding the FRT, there exists a spatial variability in the confinement along the fallopian tube^27,13^. The prolonged interaction of sperm with the surfaces of the FRT in less confined segments, including the isthmus, promotes the sperm capacitation before reaching the oocyte^22^. Higher geometric confinement in ampulla and infundibulum, zones near the fertilization site, encourages short interactions between sperm and surface but with higher frequency due to faster transit and reduced interaction duration. The reduced transition time and increased collisions maximizes the probability of reaching the oocyte’s Zona Pellucida^58^ and initiates fusion of plasma membranes^59^.

Hydrodynamic interactions between sperm and surfaces causes chirality in their motion^29,45^ restricting the diffusive behavior^60,51^. Through a mathematical model using a simple Markov model, we found up to ∼9-fold increase of diffusion in obstacle lattices compared to free swimming diffusion highlighting the role of collisions on self-diffusion of sperm^61,51^. The computational model did not predict any major difference between two lattice spacings, however, the experimentally observed peaks in the short-term direction (**Fig. 4b**) indicates a channeling effect that originates from sperm sliding along obstacles over successive collisions^43^. Stochasticity of collision and direction change especially after head-on collision ultimately randomizes the migration enabling more extensive lattice exploration. This enhanced diffusive behavior in confined lattices simulates the regions of higher confinement near the fertilization zone as sperm approach the oocyte, and the need for sperm to more widely explore a range of complex areas. Hyperactivation^55^, as a sperm motility altering mechanism near the oocyte, did not affect the overall behavior of lattice exploration and diffusive behavior.

In addition to the effect on transport behavior, sperm collisions with the circular obstacles in the device mimics the contact phase of sperm with an oocyte. We found that sperm tend to have oblique collision angles even in unconfined obstacles and in presence of the hyperactivation agent, and we suggest that sperm pre-collision chirality plays a role in preventing head-on collisions. The oblique collision angle permits the sperm to adjust the frictional contact between oocyte and sperm head maximizing the ability of the sperm to penetrate the oocyte ^62,63^. Our observation might also explain the early observations of the inclined penetration paths of sperm in the protecting layer around the oocyte known as Zona Pleucida^64^.

To conclude, how sperm interacts with and migrates within a complex environment is essential for fully describing the process of mammalian fertilization. We show how the confinement in a lattice of obstacles shape the sperm-surface interaction and migratory behavior. This is of great relevance since the fallopian tube features geometrical complexity. Ultimately, sperm migration is governed by the complexity of the surrounding environment, where the interplay of geometrical confinement, mucus viscoelasticity^45^, and biochemical signaling^65–67^ collectively influence their migratory behavior in the female reproductive tract^68^ . These factors promote topotactic guidance^68^ as sperm slide along the epithelial surface of the female reproductive tract. Future investigations should also address the influence of external cues, such as fluid flow, which can elicit rheotactic navigation^69,70^, break chirality^54^, and tune the sperm-surface interactions^25^.

## Methods

### Medium and sperm preparation

Milk-extended cryopreserved bovine semen was thawed in a water bath at 38.5 °C. The post-thawed semen sample contained 100 million/mL sperm with motility of 30%. TALP medium was prepared in sterile water as follows^42^: NaCl (110 mM), KCl (2.68 mM), NaH_2_PO_4_(0.36 mM), NaHCO_3_ (25 mM), MgCl_2_ (0.49 mM), CaCl_2_ (2.4 mM), HEPES buffer (25 mM), glucose (5.56 mM), pyruvic acid (1.0 mM), penicillin G (0.006%), and bovine serum albumin (BSA) (20 mg/mL) (pH adjusted to 7.3). BSA prevents the sperm from being tethered to the surface microfluidic walls. We used caffeine to induce hyperactivation, as it is a well-established promoter of bovine sperm hyperactivation^55^, with effects that are strongly concentration-dependent^71^.

### Microfabrication

The microfluidic device (**Fig. 1b**) was fabricated from polydimethylsiloxane (PDMS) by conventional soft lithography technique^41^. The negative photoresist SU-8 2025 (KAYAKU Advanced Materials) was coated on the wafer using spin-coating at 3000 rpm for 30 s, baked at 65 °C for 2 min and 95 °C for 5 min. The SU-8 coated wafers were exposed to 365 nm UV light through the pattern mask for 38 s and baked at 65 °C for 1 min and 95 °C for 5 min. The mold was subsequently developed in the SU-8 developer for 5 min. We also hard-baked the mold at 180 °C for 5 min to remove the cracks. The height of the fabricated structures was 35 ± 2 µm measured using a profilometer (KLA Tencor P7 profilometer). The mold was then cast with PDMS (Sylgard 184, Dow Corning) prepared according to the manufacturer’s recommended procedure (1:10 elastomer base: curing agent) and cured at 55 °C for 2 h. Inlet ports were punched, and the PDMS was plasma-bonded to a glass slide.

### Microscopy and cell tracking

The microfluidic device was maintained at 38.5 °C (bovine baseline body temperature) on a Carl Zeiss heated microscope stage in all experiments (**Fig S1**). This device was initially filled with the medium and the air bubbles removed by pressurizing the filled device using a syringe pump (Chemyx Fusion 200). Once the plugs are removed and the pressure is released a drop of medium is created on one side while the other port is plugged (**Fig S1**). A 2 µL aliquot of 4X-diluted bovine semen was added to the droplet gently and the sperm allowed to move from the droplet into the device, there was no fluid flow within the device. Before acquiring the movies, a wait period of 5 min was observed to allow the sperm to diffuse into the chamber while dead sperm remained near the deposition port. Sperm motion and their interactions with obstacles within the device were recorded using an inverted phase-contrast microscope (Nikon Eclipse TE300) equipped with a 10x objective and a digital camera (Andor Zyla 4.2). Note that sperms swimming on glass and PDMS surfaces were both tracked, however, the majority of cells were on the glass. The movies were acquired at 30 frames per second.

### Trajectory analysis

We tracked the bovine sperm while approaching and interacting with the obstacle in the standard TALP medium (**Fig. 1d**). The scattering angles were estimated by passing the sperm trajectories through a Savitzky–Golay filter^72^ (window size: 13 frames, polynomial order: 3) to smooth the paths, and then calculating the average direction of motion as the cells interacted with the obstacles. The obstacles were identified from static frames from each movie using the Hough Circle Transform^73^ with parameters adjusted to the size of the obstacles (measured manually in FIJI platform^74^) and tweaked until the expected number of obstacles was present. Using the micrometer radius of the identified obstacles as a known constant, the ratio of pixels per micrometer was estimated and used for distance unit conversions. Interactions with a given obstacle were estimated based on a drawn circle centered at the obstacle with a radius of *r_p_* + *r_s_*, where *r_p_* = 100 µm is the radius of the obstacle and *r_s_* = 9.2 µm is the length of a sperm head (**Fig. 1d**).

An interaction with an obstacle was defined as an intersection of the average path and the inner area of the drawn circle. In cases where the same smoothed trajectory appeared to have multiple interactions with the same obstacle, the interactions were joined into one if the arc length of the projection of the average path snippet between intersections onto the impact zone divided by its Euclidean distance was more than 0.9 and left as separate interactions otherwise. Collision angle (𝛼), departure angle (𝛽), and central angle (𝛾) were calculated for each scattering event. Collision and departure angles (𝛼, 𝛽 ∈[-90°,90°]) are angles between the local outward normal to surface and the incoming or outgoing swimming orientations. Central angle (𝛾) represents the angular distance travelled by sperm on surface. If 𝛾 < 0, the cell travels clockwise around the obstacle; if 𝛾 > 0, then the cell travels counterclockwise. The scattering angle calculation methods are prone to error because of the sensitivity of calculated angles on the definition of the impact zones which may generate biased data sets. Therefore, to reduce bias, we used a computational method to consider the precision in the calculation of angles by randomly rotating the trajectory of each interaction event (approach, interaction, and departure) around the detected obstacle center 50 times. The calculated standard deviation for sperm determined angles was consistently less than 0.005° indicating that any observed difference in determined parameters is either the effect of predefined geometric condition or variability in the cell population.

The swimming angle θ(t_j_) was determined by independently smoothing the x and y trajectory coordinates with a Gaussian filter (x^S^ and y^S^) to minimize high-frequency noise, and then calculating the angle using

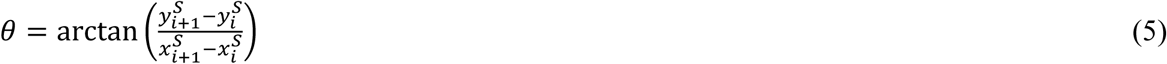

## Supplementary Information

Supporting tables, discussion and figures can be found in the supplementary information. Raw data for all figures presented in the main manuscript have been published at: https://doi.org/10.5281/zenodo.17603601 .

## Supporting information

Supplemental Figures, Tables and Discussion

## Acknowledgements

We thank Dr. James Sethna for insightful and stimulating discussions, and Dr. Kelley Donaghy for her valuable assistance in refining and editing the manuscript. We are also grateful to Dr. Adolfo Sanchez-Blanco for providing the images of the human fallopian tube cross-section. This work was performed in part at the Cornell NanoScale Facility, a member of the National Nanotechnology Coordinated Infrastructure (NNCI), which is supported by the National Science Foundation (Grant NNCI-2025233). Bovine semen samples were generously provided by Select Sires, Inc. (Plain City, OH, USA).

