## Supplemental Figures, Tables and Discussion for "Adaptive surface sensing enables mammalian sperm navigation in complex environments"

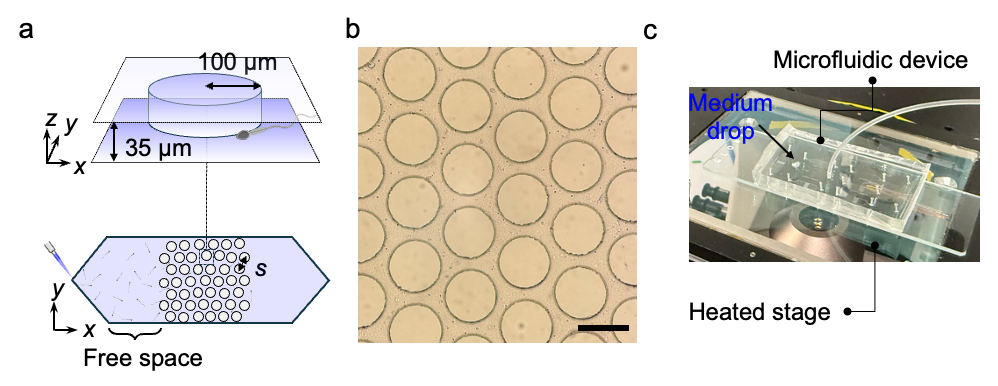

**Fig. S1** Setup used for experiments.

### Note 1. Sperm-pillar collision angle model

We consider a circle with radius of R as the pillar and a sperm following a curved trajectory with a constant curvature of 1/*r* with the circular trajectory’s center located at a distance *x* from the center of the pillar. By mirror symmetry, we can consider only the counterclockwise directions of the trajectory; by rotational invariance about the pillar center, any instance of location of the pillar and trajectory centers can be transformed to the base case shown in **Fig. S2(a)**. We also only consider cases of r>R due to our observations of the cell.

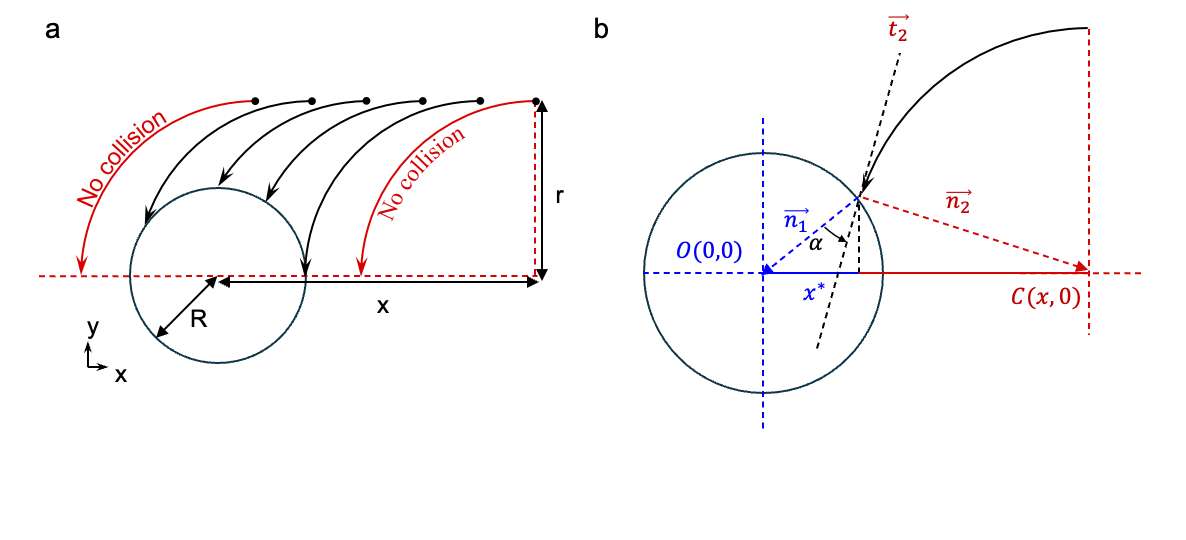

**Fig. S2.** Model setting for collision angle analysis. (a) A hypothetical sperm represented by the circle of its trajectory placed at a distance *x* from the center of the pillar travels on a curved trajectory with a radius of *r*. If *d* is out of a defined range, trajectory does not intersect the pillar. (b) Schematic of the set-up to find the collision angle $\alpha$ with depiction of all variables.

The range where the collision (intersect) exists is defined by $r-R\leq x\leq r+R$, where the bounds are the cases where the trajectory grazes the pillar surface. We expect the distribution of *x* to be uniform in space. This uniformity can be deduced from the uniformity of the collision locations on the pillar due to the randomness of the initial sperm orientations. We experimentally determined the latter by locating the collision points on the pillar surface for sperms swimming in standard medium and defining the radial location $\varphi$. The profile of probability density function (PDF) of $\varphi$ shows randomness of collision locations (**Fig. S3a**). The uniformity of the distribution is confirmed by the linearity of the cumulative density function (CDF) graph (**Fig. S3b**). This also proves the minimal effect an adjacent pillar has on its neighbors and ensures a non-biased data set for collision angles. Therefore, we define x as a continuous random variable with its probability density function (PDF) given by $P\left( x \right)=1/[2R]$.

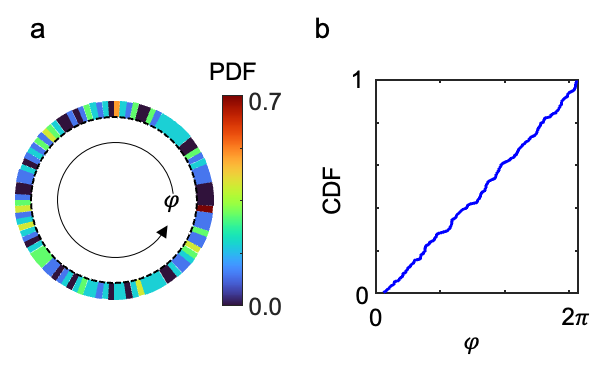

**Fig. S3**. Distribution of collision locations on the pillar for sperms swimming and colliding in the standard medium. (a) Distribution of angular location of collision $\varphi$. Color bar indicates probability density function (PDF). (b) the linearity of the cumulative density function (CDF) indicates the uniformity of the distribution.

The collision angle $\alpha$ is the angle between the inward normal vector from the center of the pillar $\vec{n_{1}}=(x^{*},y^{*})$ and the vector tangent to the sperm trajectory $\vec{t_{1}}=(y^{*},x-x^{*})$ at the collision point (x*,y*) (**Fig. S2b**). So, $\alpha$ is given by

$cos \alpha=\frac{\vec{n} .\vec{t}}{\|\vec{n}\| \|\vec{t}\|}=\frac{x y^{*}}{R r}$	(S1)

The location of the collision point $y^{*}$ is geometrically calculated using the following formula

$y^{*}=\sqrt{R^{2}-\left( \frac{R^{2}-r^{2}+x^{2}}{2x} \right)^{2}}$	(S2)

Substituting equation (1) to equation (2) gives

$cos \alpha=\frac{x}{R r}\sqrt{R^{2}-\left( \frac{R^{2}-r^{2}+x^{2}}{2x} \right)^{2}}$	(S3)

Rearranging equation (3) to get *x*

$x\left( \alpha\right)=\sqrt{R^{2}+r^{2}\pm2 R rsinsin \left( \alpha\right)}$	(S4)

Transformation of the uniform initial trajectory center distribution, $P\left( x \right)$, into the collision angle distribution is given by $P_{x}\left( \alpha\right)=P\left( x \right)\left| \frac{dx}{d\alpha} \right|$, since $\alpha(x)$ is monotonic across our range. The Jacobian of the transformation $\left| \frac{dx}{d\alpha} \right|$ is given by

$\left| \frac{dx}{d\alpha} \right|=\frac{R r coscos \left( \alpha\right)}{\sqrt{R^{2}+r^{2}\pm2R rsinsin \left( \alpha\right)}}$	(S5)

The probability density function of collision angles is then given by

$P\left( \alpha\right)=\frac{1}{2}\frac{coscos \left( \alpha\right)}{\sqrt{\left( \frac{R}{r} \right)^{2}+1\pm2\frac{R}{r}sinsin \left( \alpha\right)}}$	(S6)

Where k = R/r denotes the radius ratio. Since R<r in all conditions considered, the integral of $P\left( \alpha\right)$ over the full domain of $\alpha\in[-\frac{\pi}{2},\frac{\pi}{2}]$ evaluates to 1. Therefore, no additional normalization constant is required. This confirms that P(α) is a valid PDF describing the angular distribution of collision angles. The PDF distinguishes clockwise (CW) and counterclockwise (CCW) deflections through the “±” sign: the “+” corresponds to CCW deflections, while the “−” corresponds to CW deflections around the central axis. For comparative study, we assume only CCW collisions.

We can rewrite the k parameter in terms of the curvature of trajectory ($\kappa$) and curvature of pillar ($\kappa_{p}$): $k=\kappa/\kappa_{p}$. For straight microswimmers, $k=0$, the function simplifies to $P\left( \alpha\right)\approx\frac{1}{2}cos\left( \alpha\right)$, which is similar to the distribution function derived by Jakuszeit et al^2^. This explains that the most likely collision angles are head-on collisions ($\alpha=0$) and grazing events ($\alpha=\pm\frac{\pi}{2}$) are rare. The distribution of collision angles with a constant pillar radius for curved trajectories depends on k. Comparing the distribution for different ratios of k, we observed that increasing the intrinsic curvature of swimmer rotation shifts the distribution toward non-zero medians. A special case is when r = R ($k=1$) which indicates that the most probable collision angle is close to $\frac{\pi}{2}$.

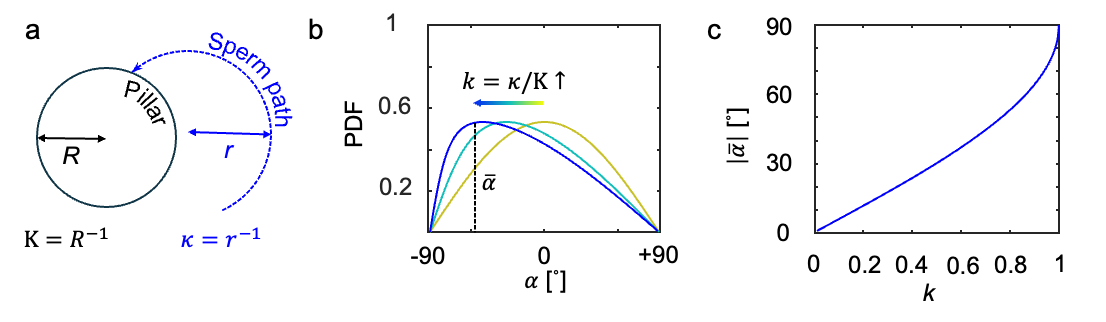

**Fig. S4** Effect of chirality of moving particles on collision angle distribution.

### Note 2. Interaction length of sperm on obstacle surface

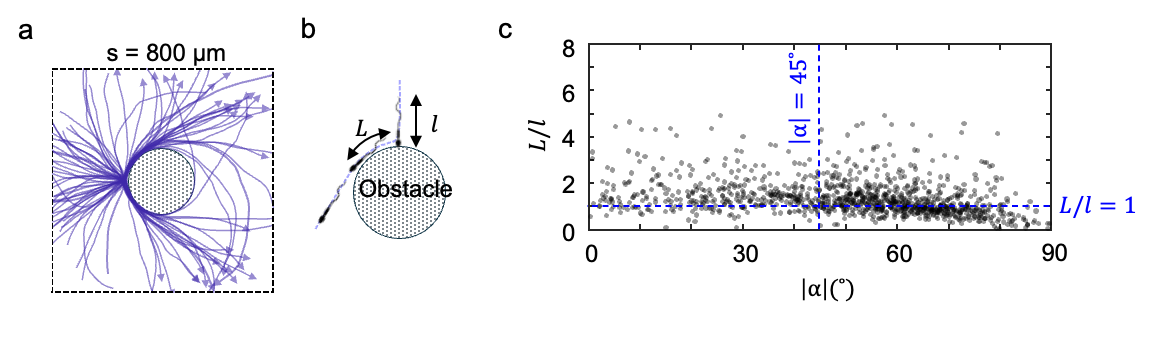

**Fig. S5** Interaction length. (a) Trajectories of sperm interacting with an unconfined obstacle. Arrows show the direction of swimming. (b) Schematic indicating the parameters. (c) Majority of cells have an interaction length in the order of sperm body length (*l* = 60 µm).

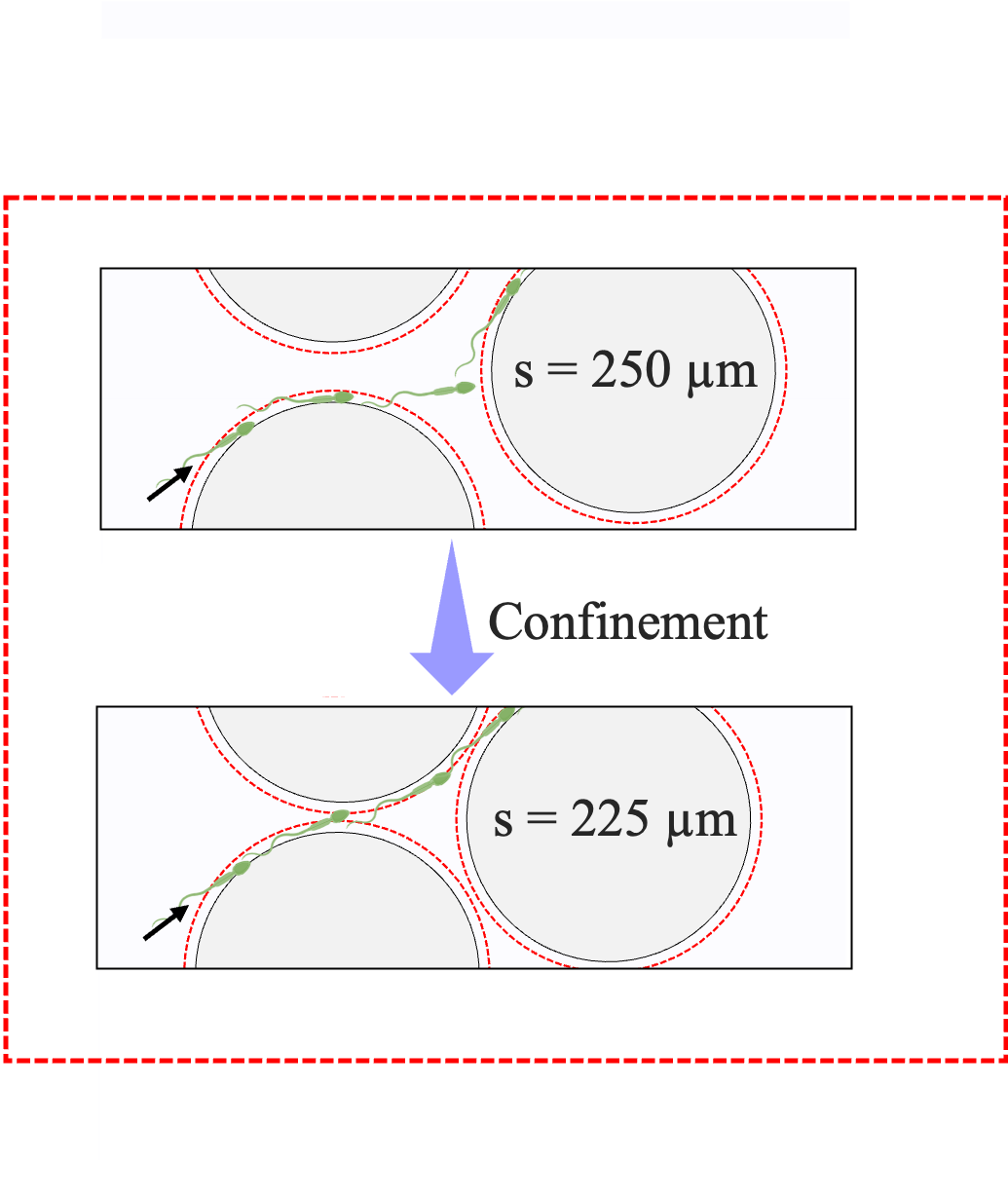

**Fig. S6** Reduced interaction time in more confined lattice due to near field effects of neibor obstacle.

### Note 3. Effective diffusivity

Calculation of D_eff_:

We consider the two-state dynamics for sperm migrating in a lattice of obstacles (see main text for details). Integrating the full space-dependent master equations (equation (1) in main text) yields

$\frac{\partial p_{1}}{\partial t}=-k_{D}p_{1}+k_{C}p_{0}$	(S7 a]

$\frac{\partial p_{0}}{\partial t}=-V_{0}\nabla\left( \hat{e}\left( \theta\right)p_{0} \right)-\Omega_{0}\partial_{\theta}p_{0}+D_{\theta}\partial_{\theta\theta}p_{0}-k_{C}p_{0}+k_{D}\int g\left( \theta-\theta^{'} \right)p_{0}(\theta^{'})d\theta'$	(S7 b)

The interaction (stop) state simply mediates between transits (runs), while sperm undergo rotation and angular diffusion during the transit.

Applying Fourier expansion

$p_{i}\left( \theta,t \right)=\sum_{n} \hat{p}_{i}\left( n,t \right)e^{in\theta}, \hat{g}\left( n \right)=\int e^{in\psi}g\left( \psi\right)d\psi$	(S8)

the angular convolution reduces to multiplication. For each mode *n* we have,

$\frac{\partial\hat{p}_{1}\left( n,t \right)}{\partial t}=-k_{D}\hat{p}_{1}\left( n,t \right)+k_{C}\hat{p}_{0}\left( n,t \right)$	(S9 a)

$\frac{\partial\hat{p}_{0}\left( n,t \right)}{\partial t}=\left( in\Omega_{0}-k_{C}-n^{2}D_{\theta} \right)\hat{p}_{0}\left( n,t \right)-k_{D}\hat{g}\left( n \right)\hat{p}_{1}\left( n,t \right)$	(S9 b)

Since translational transport couples only to the first Fourier mode, we set n = 1. Defining

$p_{i}\left( t \right)\equiv\hat{p}_{i}\left( 1,t \right), G\equiv\hat{g}\left( 1 \right)=\langle coscos \psi\rangle$	(S10)

The system then reads

$\frac{\partial}{\partial t}\left( p_{1} p_{0} \right)=M\left( p_{1} p_{0} \right), M=\left( -k_{D} k_{C} k_{D}G -(k_{C}+D_{\theta}+i\Omega_{0}) \right)$	(S11)

With Laplace transform $P_{i}\left( s \right)=\int_{0}^{\infty} e^{-st}p_{i}\left( t \right)dt$and initial condition corresponding to a run (transit state) at angle 0,

$p_{0}\left( 0 \right)=\frac{1}{2\pi}, p_{1}\left( 0 \right)=0$	(S12)

the algebraic system yields

$P_{0}\left( s \right)=\frac{s+k_{D}}{D\left( s \right)}.\frac{1}{2\pi}, D\left( s \right)=\left( s+k_{D} \right)\left( s+k_{C}+D_{\theta}+i\Omega_{0} \right)-k_{C}k_{D}G.$	(S13)

At s=0,

$P_{0}\left( 0 \right)=\frac{1}{2\pi}\frac{1}{D_{\theta}+k_{C}\left( 1-G \right)+i\Omega_{0}}$	(S14)

The velocity autocorrelation function is

$C_{v}\left( t \right)=\langle v\left( 0 \right).v\left( t \right)\rangle=2\pi V_{0}^{2}p_{0}^{eq}Re\left\{ p_{0}\left( t \right) \right\}$	(S15)

where the steady-state transit fraction is

$p_{0}^{eq}=\frac{k_{D}}{k_{C}+k_{D}}$	(S16)

Applying Taylor-Kubo relation,

$D_{eff}=\frac{1}{2}\int_{0}^{\infty} C_{v}\left( t \right) dt=\pi V_{0}^{2}p_{0}^{eq}Re\{P_{0}(0)\}$	(S17)

Gives the analytical expression for the effective diffusivity

$D_{eff}=\frac{V_{0}^{2}}{2}\frac{k_{D}}{k_{C}+k_{D}}\frac{D_{\theta}+k_{C}\left( 1-G \right)}{\left[ D_{\theta}+k_{C}\left( 1-G \right) \right]^{2}+\Omega_{0}^{2}}$	(S18)

Experimental measurements of $\psi$ indicated a Gaussian distribution (**Fig. 2c** in main text). For a Gaussian kernel g($\psi$) of variance $\sigma^{2}$,

$G=\langle coscos \psi\rangle=e^{-\frac{\sigma^{2}}{2}}$,	(S19)

so the diffusivity reduces to

$D_{eff}=\frac{V_{0}^{2}}{2}\frac{k_{D}}{k_{C}+k_{D}}\frac{D_{\theta}+k_{C}\left( 1-e^{-\frac{\sigma^{2}}{2}} \right)}{\left[ D_{\theta}+k_{C}\left( 1-e^{-\frac{\sigma^{2}}{2}} \right) \right]^{2}+\Omega_{0}^{2}}$	(S20)

Some limiting behaviors include:

1) No stops (k_C_=0, free space swimming):

$D_{eff}=\frac{V_{0}^{2}}{2}\frac{D_{\theta}}{D_{\theta}^{2}+\Omega_{0}^{2}}$	(S21)

Which is the standard effective diffusivity for a chiral active particle.

2) Stop-dominated reorientation ($D_{\theta}\ll k_{C}(1-G)$):

$D_{eff}\approx D_{\theta}+\frac{k_{C}\sigma^{2}}{2}$	(S22)

3) Strong chirality limit ($|\Omega_{0}|\gg D_{\theta}+k_{C}(1-G)$):

$D_{eff}=\frac{V_{0}^{2}}{2}\frac{k_{D}}{k_{C}+k_{D}}\frac{D_{\theta}+k_{C}\left( 1-e^{-\frac{\sigma^{2}}{2}} \right)}{D_{\theta}^{2}+\Omega_{0}^{2}}$	(S23)

which suppresses the long-term translational diffusion.

Measurement of *V_0_*, *𝛺_0_*, and D_θ_ in free space:

The instantaneous linear velocity v(t) of each tracked cell was computed from the trajectory positions (x_i_,y_i_), recorded at discrete time points $t_{i}=i\Delta t$, where $\Delta t$ =33.3 ms is the frame interval:

$v_{i}=\frac{\sqrt{\left( x_{i+1}-x_{i} \right)^{2}+\left( y_{i+1}-y_{i} \right)^{2}}}{\Delta t}$	(S24)

where x_i_ and y_i_ are Cartesian coordinates of the cell at frame *i*. For each trajectory (**Fig. S7a**) with N number of frames, the mean linear velocity V_0_ was calculated as

$V_{0}=\frac{1}{N-1}\sum_{i=1}^{N-1} v_{i}$	(S25)

Note that we averaged each point over 6 frames to remove the frame-to-frame noise.

Sperm exhibit systematic rotation (chirality). The mean angular velocity $\Omega_{0}$ was calculated by

$\Omega_{0}=\frac{1}{N-1} \sum_{i=1}^{N-1} \frac{\theta_{i+1}-\theta_{i}}{\Delta t}$	(S26)

where $\theta_{i}$ is the orientation of the cell at frame *i* (**Fig. S7b**). Equivalent radius of rotation can be defined as r = V_0_/$\Omega_{0}$. For each time lag t, the angular displacement relative to the mean rotation was calculated
$\Delta\theta_{i}\left( t \right)=\theta_{i+t}-\theta_{i}-\Omega_{0}t$	(S27)

mean squared angular displacement (MSAD) was computed as

$MSAD\left( t \right)=\frac{1}{N-t}\sum_{i=1}^{N-t} \left( \Delta\theta_{i}\left( t \right) \right)^{2}$	(S28)

Note that we subtracted systematic rotational diffusion ($\Omega_{0}t$) to obtain the stochastic rotational diffusion. For a purely diffusive angular motion, MSAD scales linearly with lag time *t*

$MSAD\left( t \right)\approx2D_{\theta}t$	(S29)

Therefore, the slope of the MSAD-t plot gives the rotational diffusion coefficient (**Fig. S7c**)

$D_{\theta}=\frac{1}{2}\frac{dMSAD}{dt}$	(S30)

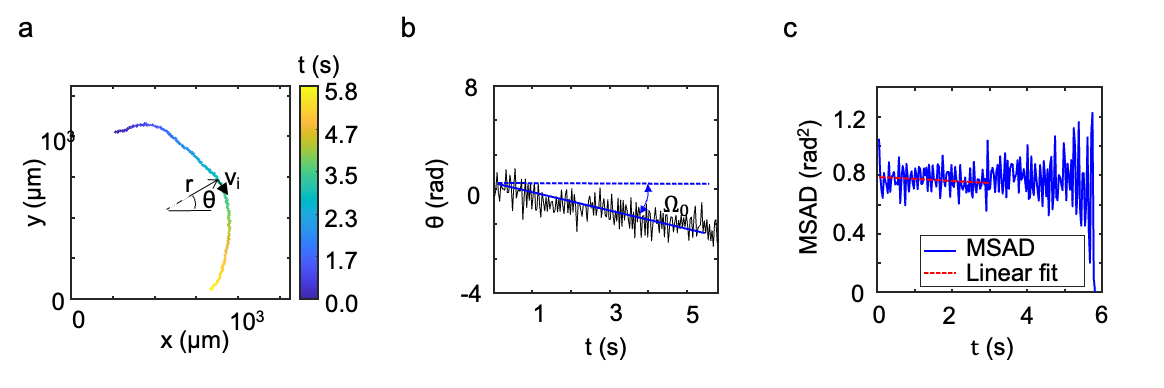

**Fig. S7** Calculation of free-swimming motility parameters. (a) Example sperm trajectory color coded with respect to time progression. *r* is the equivalent radius of rotation. (b) Instantaneous direction $\theta$ of sperm over time *t*. (c) Mean squared angular displacement (MSAD) over lag time *t*.

### Note 4. Hyperactivated sperm

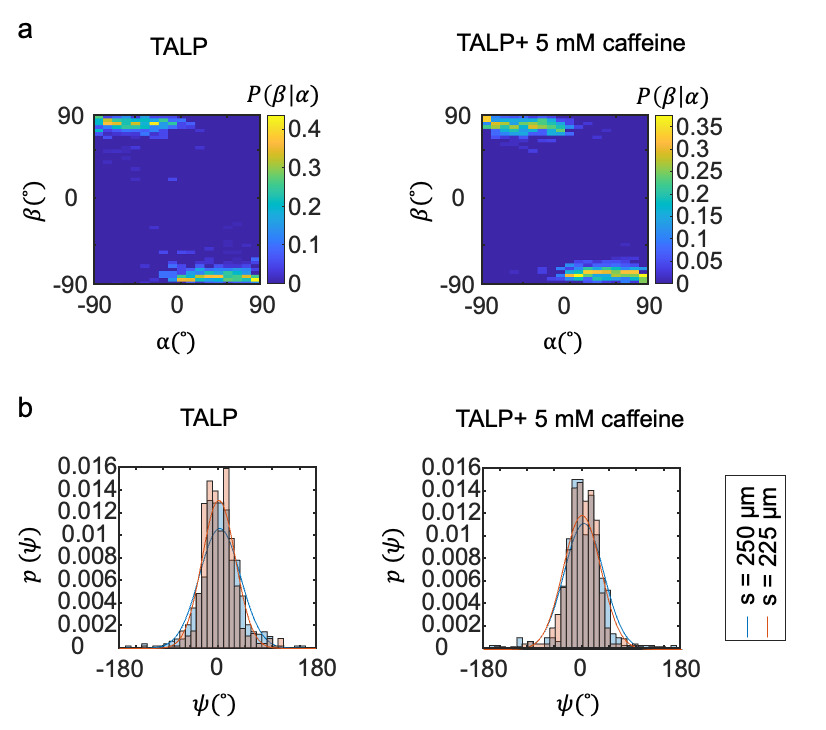

**Fig. S8** Scattering behavior of hyperactivated sperm. (a) Comparing the 𝛼–𝛽 conditional probability maps for normal (TALP) and hyperactivated (TALP+ 5mM caffeine) cells. (b) Distributions of net reorientation 𝜓 for two conditions. Solid lines are the fit to the Gaussian function.

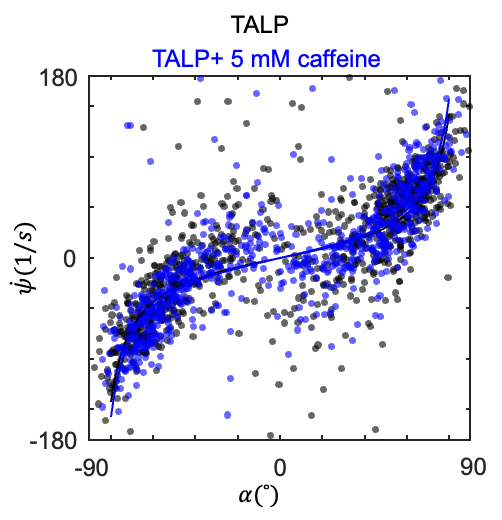

**Fig. S9** Correlation of net reorientation rate 𝜓^. with collision angle 𝛼 for normal and hyperactivated sperm. Solids lines are fitted curves to function $\dot{\psi}$= k tan (𝛼). k value is similar for both conditions. Plot is compiled from all scattering data from lattice spacings *s* = 250 µm and *s* = 225 µm.

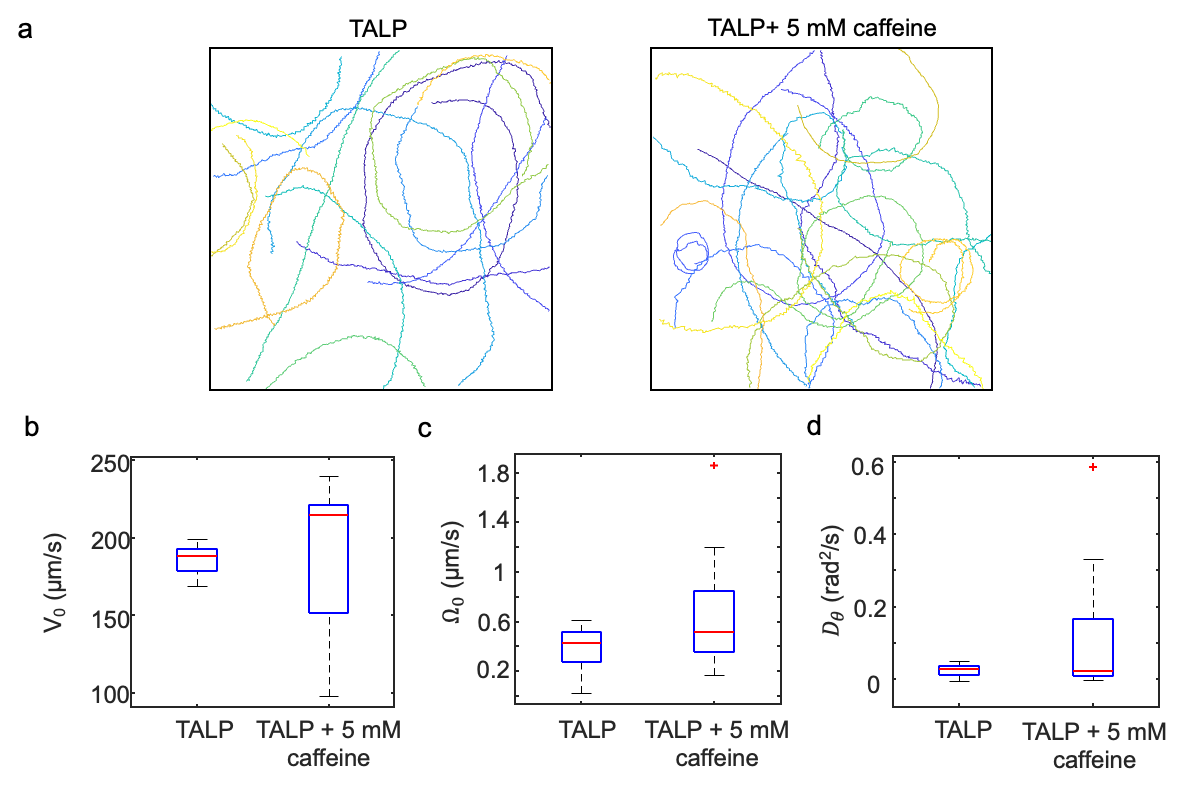

**Fig. S10** Comparing free-space motility of normal (TALP) and hyperactivated (TALP + 5 mM caffeine). (a) Representative trajectories, (b) mean linear velocity, (c) mean angular velocity, and (d) angular diffusion coefficient for each condition.

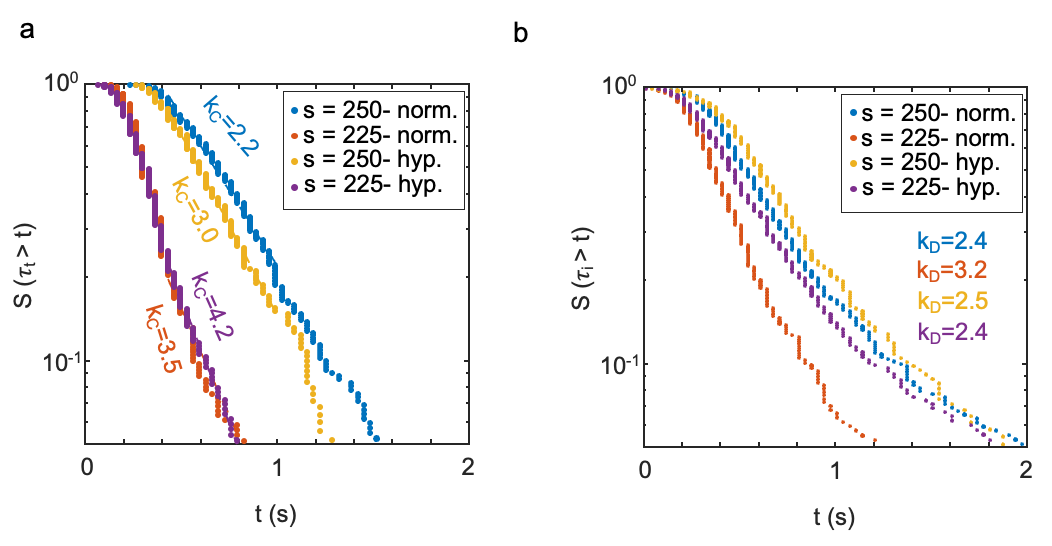

**Fig. S11** Comparing the survival curves of (a) transit duration, and (b) interaction duration for normal and hyperactivated sperm in 250 µm and 225 µm lattices. k_C_ and k_D_ are obtained from the slopes.
